## Supplemental Information for "Anionic Phospholipids Control Mechanisms of GPCR-G Protein Recognition"

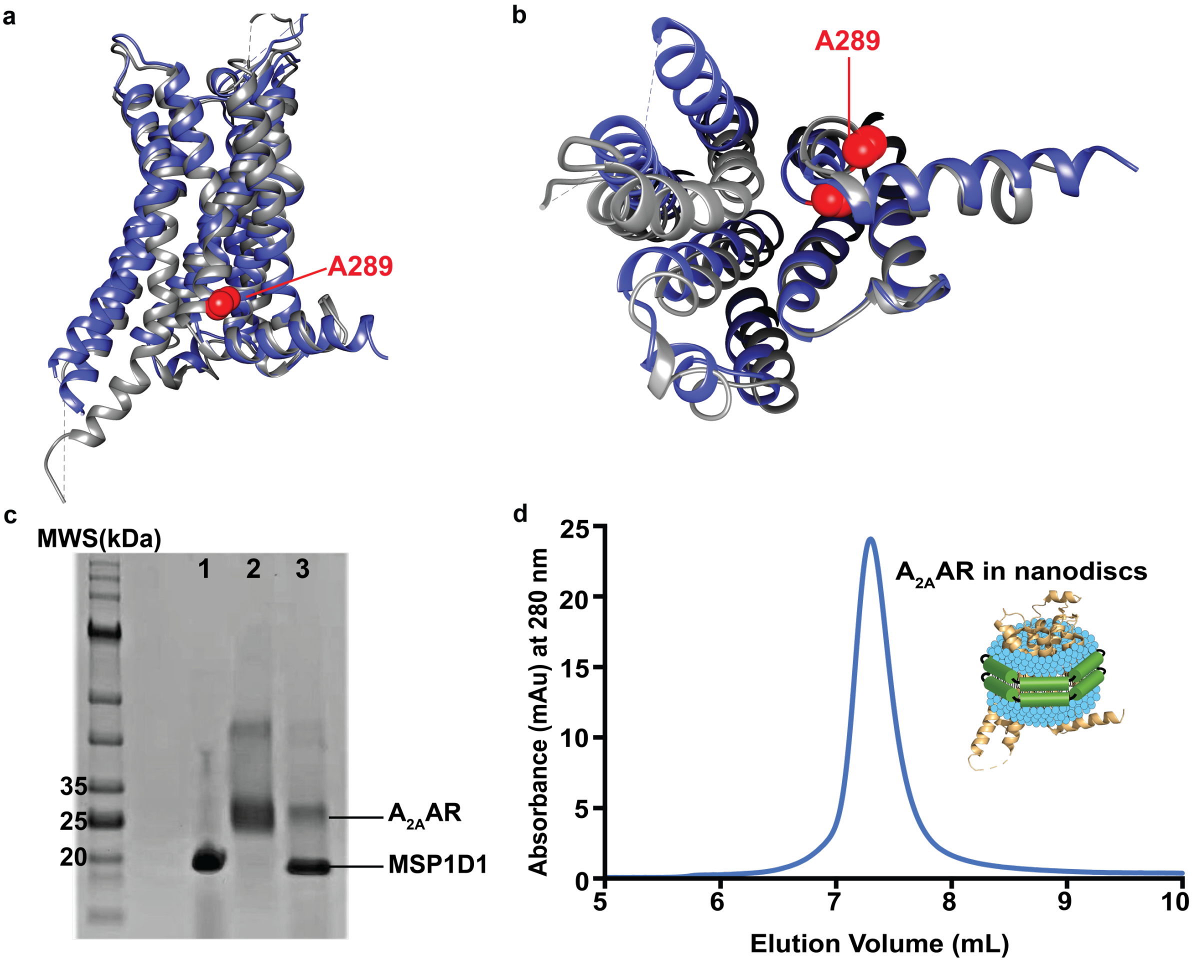


**Supplementary Fig. 1. Location of A289 in $A_{2A}AR$ crystal structures and characterization of $A_{2A}AR[A289C^{TET}]$ in lipid nanodiscs. a and b** Superposition of crystal structures of $A_{2A}AR$ in complex with the antagonist ZM241385 (grey, PDB 3EML)[1] and in a tertiary complex with the agonist NECA and mini-G$\alpha_s$ protein (blue, PDB 5G53)[2] shown in ribbon representation. The location of A289 in helix VII is represented by red spheres. **c** Representative SDS-PAGE gel showing purified MSP1D1 (lane 1), purified $A_{2A}AR[A298C^{TET}]$ in DDM/CHS mixed micelles (lane 2), and purified lipid nanodiscs containing $A_{2A}AR[A298C^{TET}]$ (lane 3). The bands corresponding to $A_{2A}AR[A298C^{TET}]$ and MSP1D1 are annotated. The lane labelled "MWS" corresponds to molecular weight standards. **d** Representative analytical size exclusion chromatogram of antagonist-bound $A_{2A}AR[A289C^{TET}]$ in lipid nanodiscs containing a mixture of POPC and POPS in a 70:30 molar ratio.

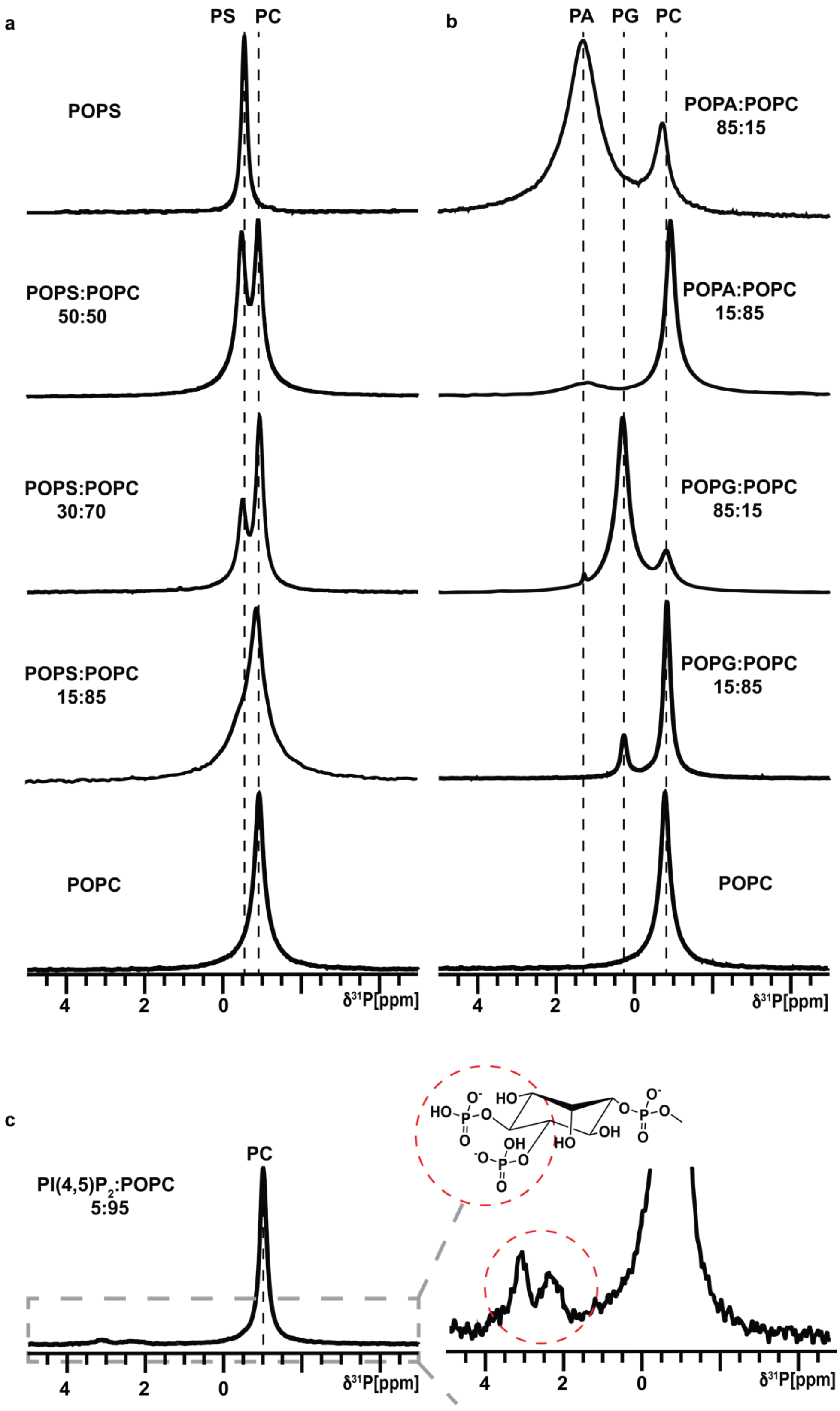

a
PS
PC
POPS
POPS:POPC
50:50
POPS:POPC
30:70
POPS:POPC
15:85
POPC
4
2
0
δ31P[ppm]
b
PA
PG
PC
POPA:POPC
85:15
POPA:POPC
15:85
POPG:POPC
85:15
POPG:POPC
15:85
POPC
4
2
0
δ31P[ppm]
c
PI(4,5)P2:POPC
5:95
PC
4
2
0
δ31P[ppm]
4
2
0
δ31P[ppm]

**Supplementary Fig. 2. Validation of nanodisc lipid composition with $^{31}$P-NMR in aqueous solutions.** **a** The 1D $^{31}$P-NMR spectra of $A_{2A}$AR[A289C$^{TET}$] in nanodiscs containing mixtures of POPC and POPS at different molar ratios, as indicated. The vertical dashed lines indicate the observed chemical shifts for the two lipid headgroups, which were consistent with manufacturer's reported values. The relative intensities of the two observed signals were quantitatively consistent with the expected lipid ratios. **b** The 1D $^{31}$P-NMR spectra of $A_{2A}$AR[A289C$^{TET}$] in nanodiscs containing mixtures of POPC with the anionic lipids POPA and POPG at different molar ratios. Same presentation details as in **a**. **c** The 1D $^{31}$P-NMR spectra of nanodiscs containing the anionic lipid PI(4,5)$P_2$ and POPC in a molar ratio of 5:95. The right panel is an expanded view of the region in the grey dashed box shown on the left, highlighting signals from the two phosphates of the inositol group.

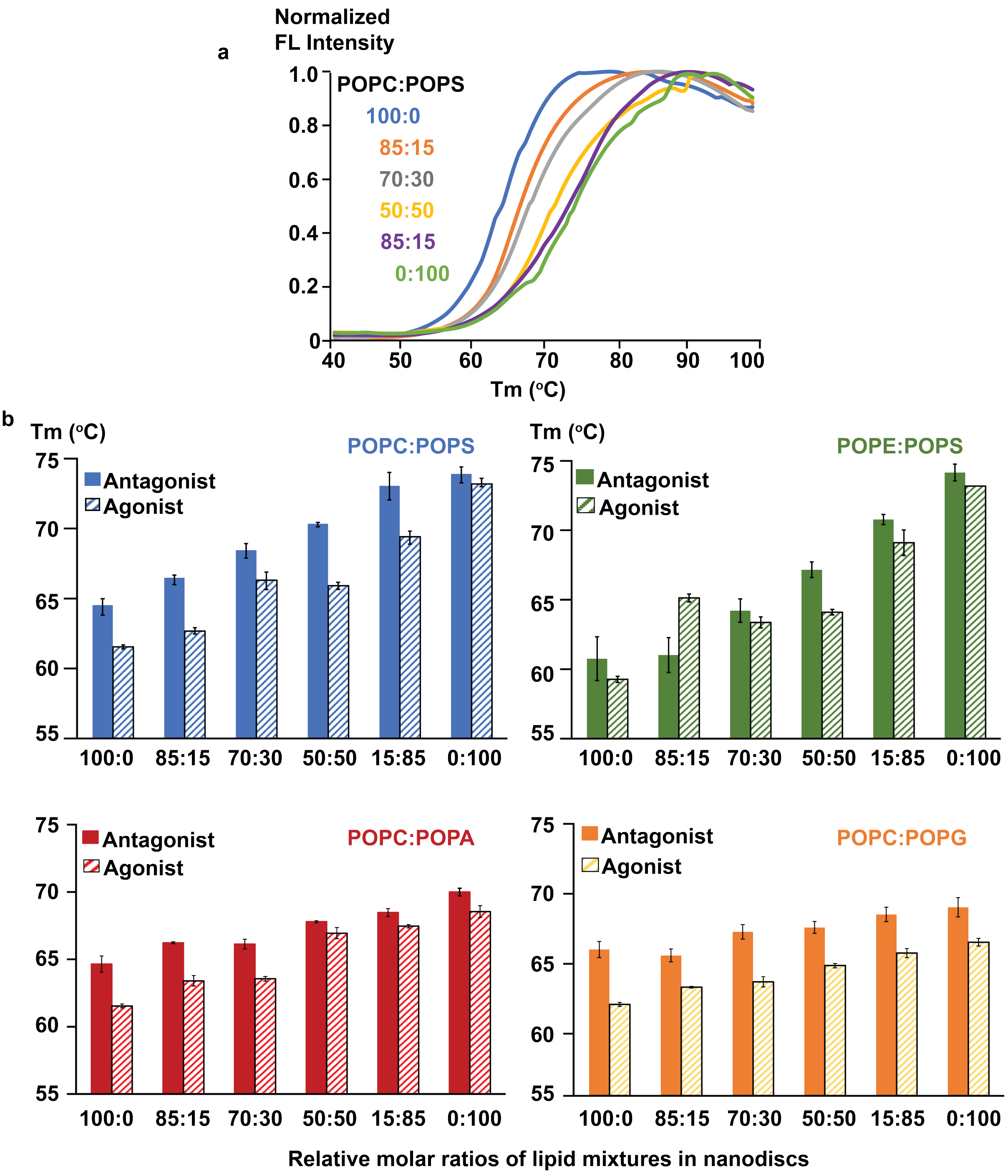
a
Normalized
FL Intensity
POPC:POPS
100:0
85:15
70:30
50:50
85:15
0:100
1.0
0.8
0.6
0.4
0.2
0
40
50
60
70
80
90
100
Tm (°C)
b
Tm (°C)
POPC:POPS
Antagonist
Agonist
Tm (°C)
POPE:POPS
Antagonist
Agonist
POPC:POPA
Antagonist
Agonist
POPC:POPG
Antagonist
Agonist
75
70
65
60
55
100:0
85:15
70:30
50:50
15:85
0:100
Relative molar ratios of lipid mixtures in nanodiscs

**Supplementary Fig. 3. Fluorescence thermal shift data of $A_{2A}$AR in nanodiscs of varying binary lipid compositions. a** Representative fluorescence thermal shift data for $A_{2A}$AR[A289C] in complex with the antagonist ZM241385 in nanodiscs containing different ratios of POPC and POPS. **b** Melting temperatures determined from fitting fluorescence thermal shift data with $A_{2A}$AR[A289C] in nanodiscs containing variable mixtures of different zwitterionic lipids (POPC and POPE) and anionic lipids (POPS, POPA, and POPG). Melting temperatures are shown for both $A_{2A}$AR[A289C] in complex with the antagonist ZM241385 (solid bars) and the agonist NECA (striped bars) for all lipid compositions. Error bars for each sample were calculated as the standard error of mean (s.e.m) for n>3 independent experiments.

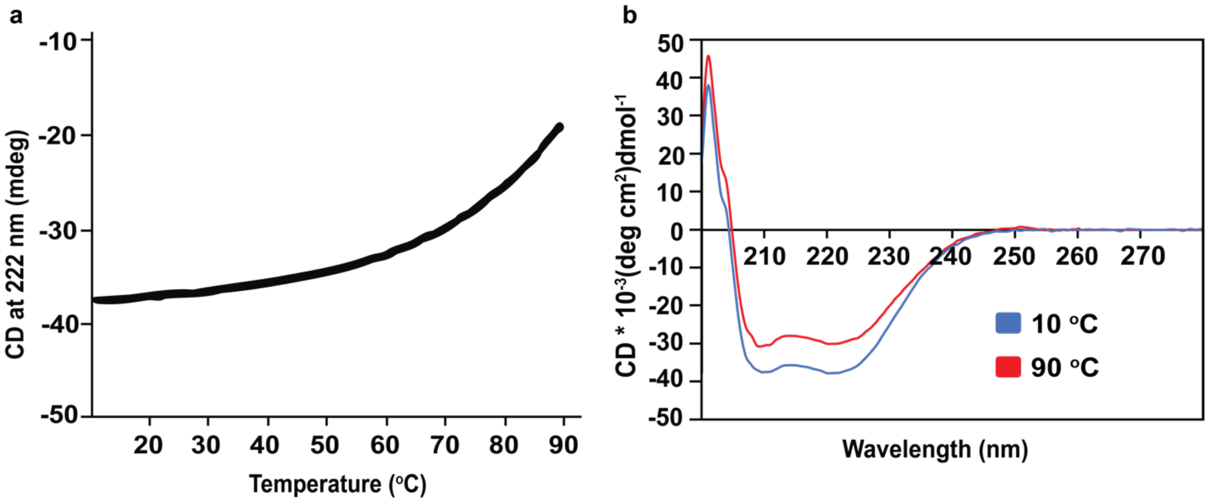


**Supplementary Fig. 4. Circular dichroism analysis of lipid nanodiscs without receptors.** **a** Variable temperature CD absorption spectrum monitored at 222 nm for nanodiscs without receptors and containing POPC and POPS at a molar ratio of 70:30. **b** CD absorption spectra of nanodiscs without receptor containing POPC and POPS at a molar ratio of 70:30 recorded at 10 °C (blue) and 90 °C (red).

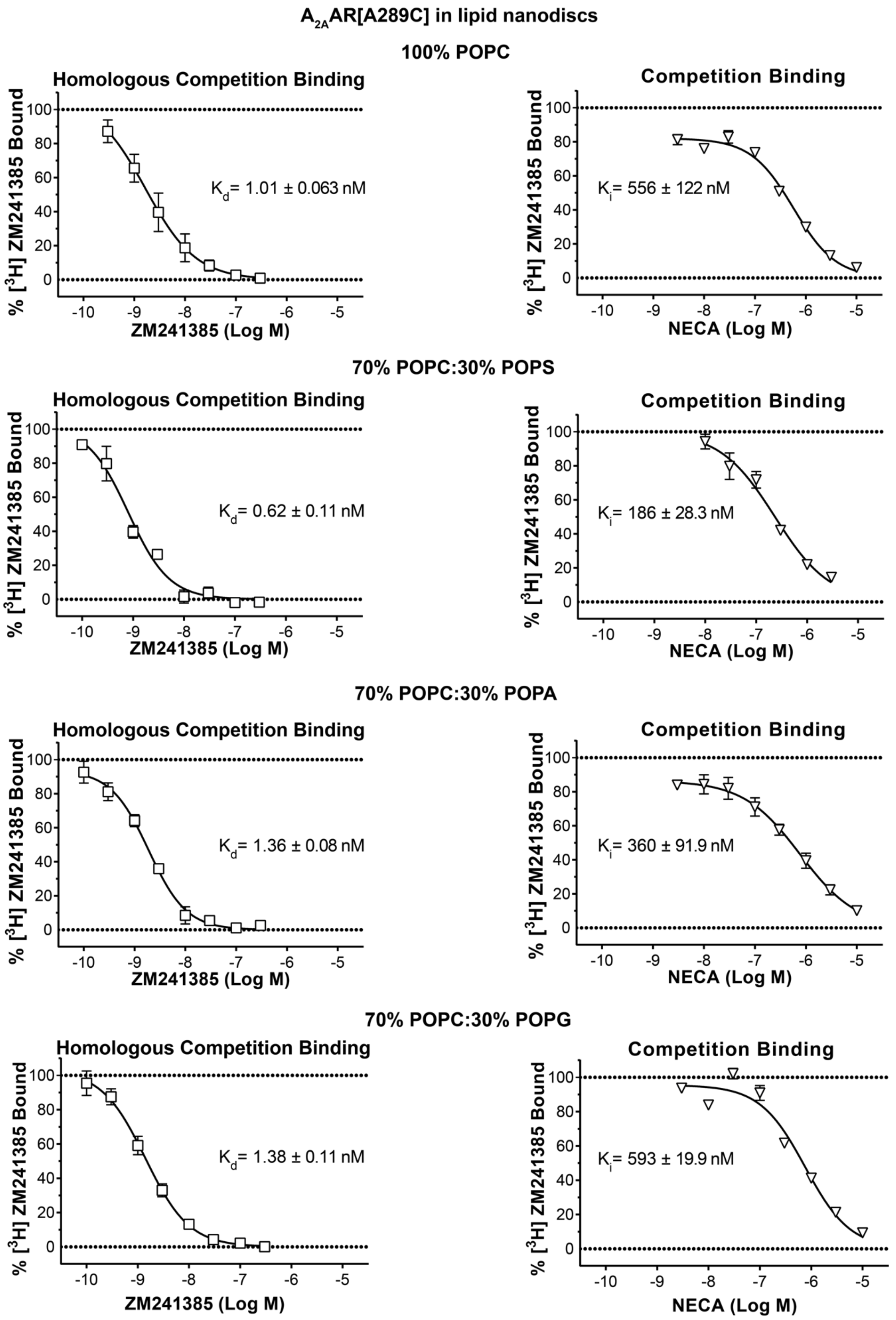

A2AAR[A289C] in lipid nanodiscs
100% POPC
Homologous Competition Binding
Kd= 1.01 ± 0.063 nM
% [3H] ZM241385 Bound
ZM241385 (Log M)
Competition Binding
Ki= 556 ± 122 nM
NECA (Log M)
70% POPC:30% POPS
Homologous Competition Binding
Kd= 0.62 ± 0.11 nM
Competition Binding
Ki= 186 ± 28.3 nM
70% POPC:30% POPA
Homologous Competition Binding
Kd= 1.36 ± 0.08 nM
Competition Binding
Ki= 360 ± 91.9 nM
70% POPC:30% POPG
Homologous Competition Binding
Kd= 1.38 ± 0.11 nM
Competition Binding
Ki= 593 ± 19.9 nM

**Supplementary Fig. 5. Pharmacological activity of $A_{2A}AR$[A289C] in lipid nanodiscs containing different lipid compositions.** Homologous competition binding experiments with the antagonist ZM241385 are shown in the left panels, and competition binding experiments with the agonist NECA are shown in the right panels. Data were measured for $A_{2A}AR$[A289C] prepared in nanodiscs containing different lipid compositions, as indicated above the panels. The measured $K_D$ or $K_I$ values are shown in each panel. Error bars indicate the s.e.m for 3 independent trials done in triplicate.

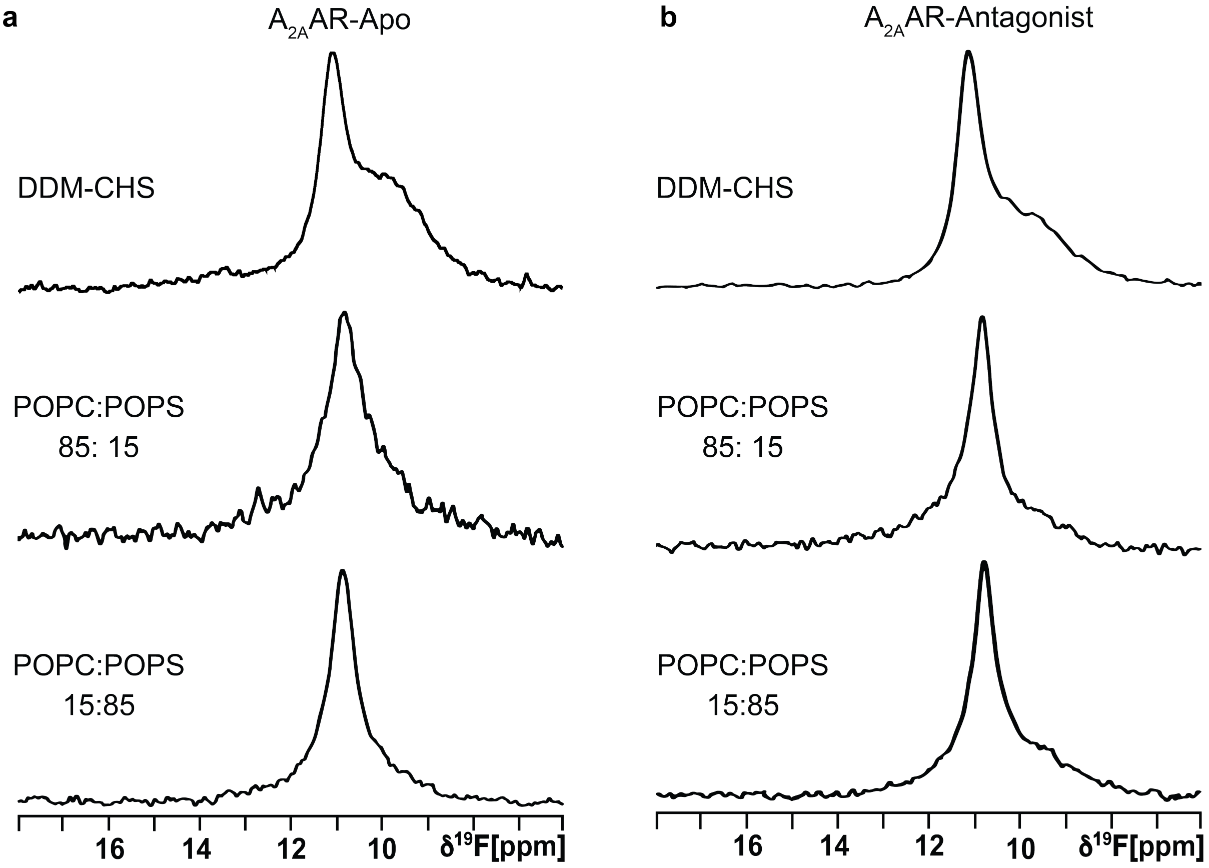


**Supplementary Fig. 6. $^{19}$F-NMR spectra of $A_{2A}$AR[A289C$^{TET}$] with no ligand added (apo) and in complex with the antagonist ZM241385 for different membrane mimetic environments. a** $^{19}$F-NMR spectra of apo $A_{2A}$AR[A289C$^{TET}$] in DDM/CHS mixed micelles and in nanodiscs composed of two different molar ratios of POPC and POPS, as indicated. **b** $^{19}$F-NMR spectra of the $A_{2A}$AR[A289C$^{TET}$] complex with the antagonist ZM241385 in DDM/CHS mixed micelles and in nanodiscs composed of two different molar ratios of POPC and POPS, as indicated.

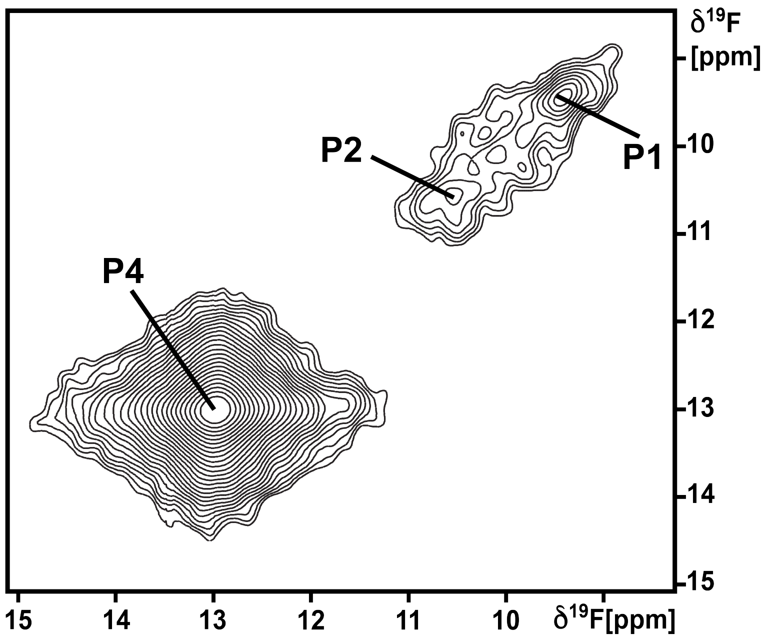


**Supplementary Fig. 7. 2D [$^{19}$F,$^{19}$F]-EXSY spectrum of $A_{2A}$AR in lipid nanodiscs.** Conformational exchange in the $A_{2A}$AR[A289C$^{TET}$] complex with the full agonist NECA in lipid nanodiscs composed of POPC and POPS in a 15:85 molar ratio observed by 2-dimensional exchange spectroscopy (EXSY). A contour plot of the 2D [$^{19}$F, $^{19}$F]-EXSY spectrum is shown, recorded at 280 K with a mixing time of 100 ms. The diagonal peak positions of P1, P2 and P4 are labelled.

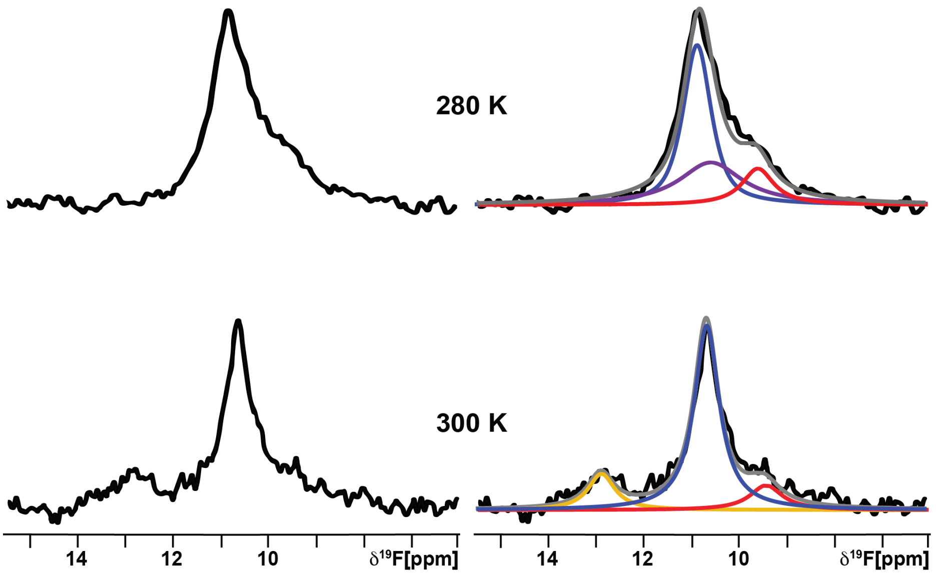


**Supplementary Fig. 8. Variable temperature $^{19}$F-NMR spectra of $A_{2A}$AR[A289C$^{TET}$] in complex with the agonist NECA in nanodiscs containing 100% POPC.** $^{19}$F-NMR spectra of the $A_{2A}$AR[A289C$^{TET}$] complex with the agonist NECA in nanodiscs containing 100% POPC measured at 280 K (top) and 300 K (bottom). Other presentation details are the same as Figure 1.

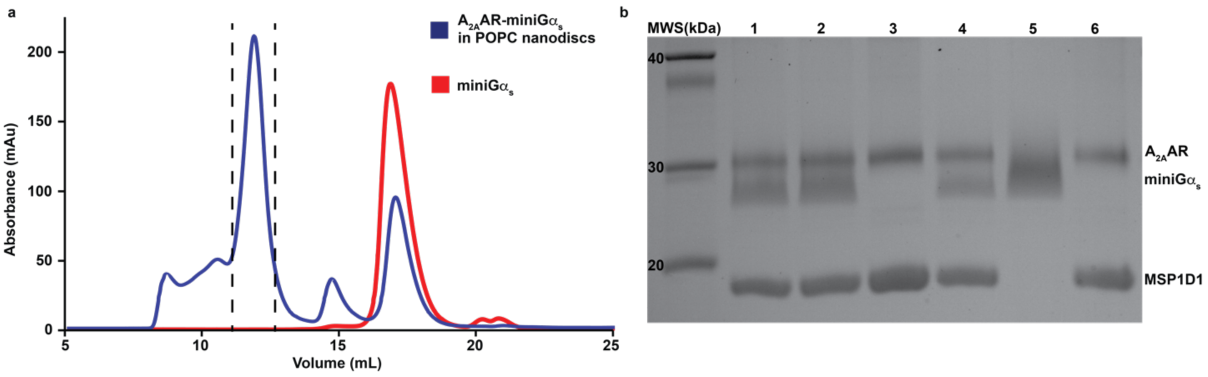


**Supplementary Fig. 9. SEC and SDS-PAGE characterization of $A_{2A}$AR-miniG$\alpha_s$ complexes in nanodiscs. a** Analytical size exclusion chromatogram of a tertiary complex of $A_{2A}$AR[A289C$^{TET}$] with the agonist NECA and miniG$\alpha_s$ in nanodiscs containing POPC lipids (blue). A chromatogram of just miniG$\alpha_s$ is superimposed and shown in red. The vertical dashed lines correspond to collected fractions pooled and run in lane 4 on the corresponding SDS-PAGE gel. **b** SDS-PAGE gel with lanes containing the following: lanes 1 and 2: purified $A_{2A}$AR[A289C$^{TET}$] in complex with the agonist NECA and miniG$\alpha_s$ in POPC:POPS (70:30) nanodiscs; lane 3: $A_{2A}$AR[A289C$^{TET}$] in complex with NECA in POPC:POPS (70:30) nanodiscs; lane 4: fractions containing the $A_{2A}$AR[A289C$^{TET}$] complex with NECA and miniG$\alpha$S in POPC nanodiscs pooled from the region of the aSEC indicated by the vertical dashed lines, lane 5: $A_{2A}$AR[A289C$^{TET}$] in complex with NECA and miniG$\alpha$S in DDM/CHS mixed micelles, and lane 6: same sample as lane 3. The bands corresponding to $A_{2A}$AR, miniG$\alpha_s$, and MSP1D1 are annotated. The lane labelled "MWS" corresponds to molecular weight standards.

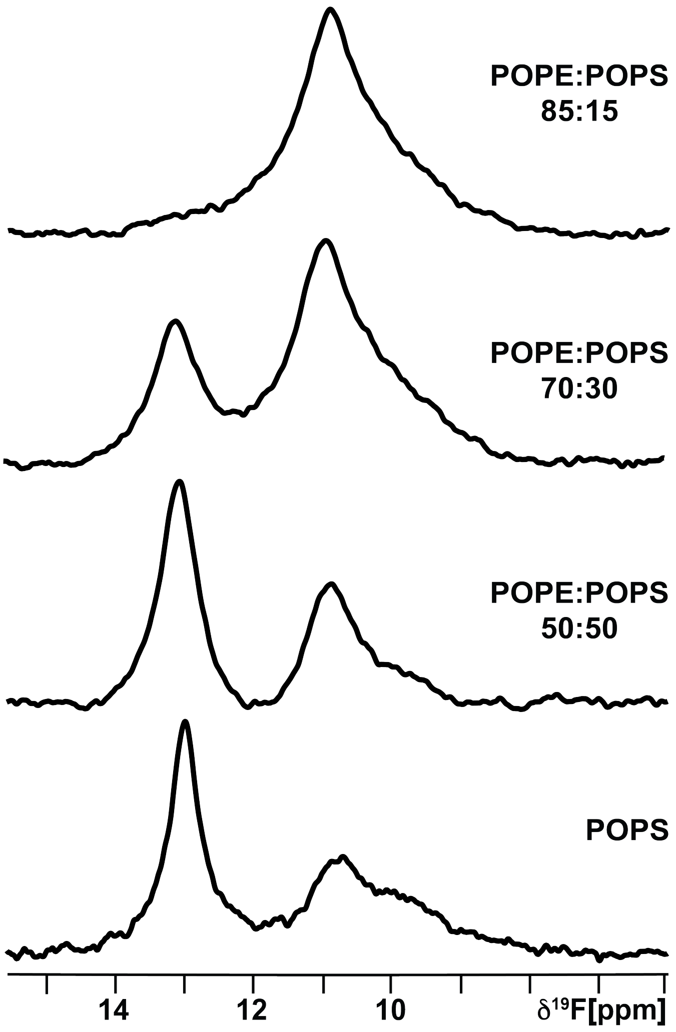


**Supplementary Fig. 10. $^{19}$F-NMR spectra of A$_{2A}$AR[A289C$^{TET}$] in complex with the agonist NECA in nanodiscs containing mixtures of POPE and POPS.** $^{19}$F-NMR spectra of the A$_{2A}$AR[A289C$^{TET}$] complex with the agonist NECA in nanodiscs containing variable amounts of the zwitterionic lipid POPE and anionic lipid POPS, as indicated in each spectrum.

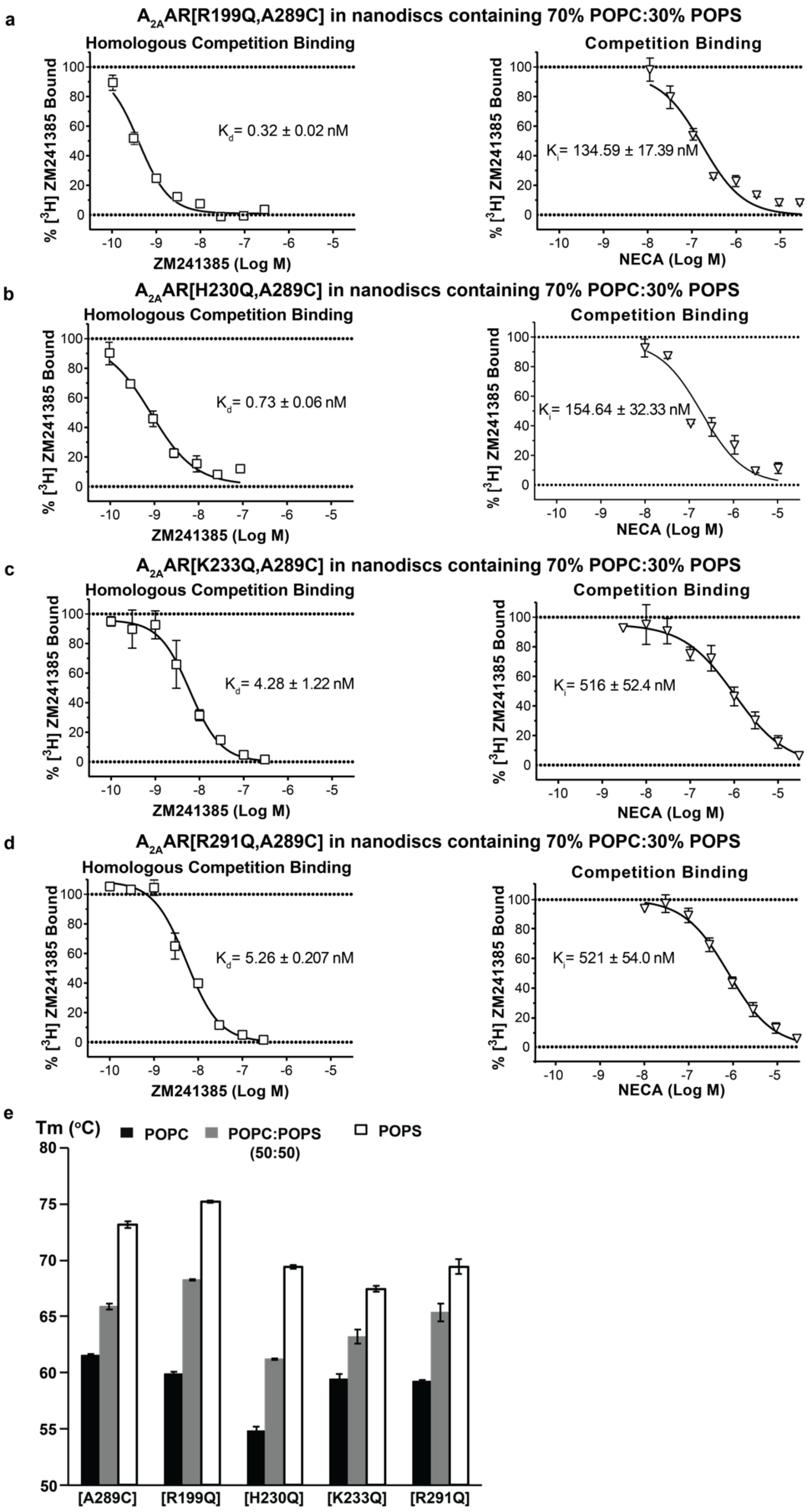
a
A2AAR[R199Q,A289C] in nanodiscs containing 70% POPC:30% POPS
Homologous Competition Binding
Kd= 0.32 ± 0.02 nM
% [3H] ZM241385 Bound
ZM241385 (Log M)
Competition Binding
Ki= 134.59 ± 17.39 nM
NECA (Log M)
b
A2AAR[H230Q,A289C] in nanodiscs containing 70% POPC:30% POPS
Homologous Competition Binding
Kd= 0.73 ± 0.06 nM
Competition Binding
Ki= 154.64 ± 32.33 nM
c
A2AAR[K233Q,A289C] in nanodiscs containing 70% POPC:30% POPS
Homologous Competition Binding
Kd= 4.28 ± 1.22 nM
Competition Binding
Ki= 516 ± 52.4 nM
d
A2AAR[R291Q,A289C] in nanodiscs containing 70% POPC:30% POPS
Homologous Competition Binding
Kd= 5.26 ± 0.207 nM
Competition Binding
Ki= 521 ± 54.0 nM
e
Tm (°C)
POPC
POPC:POPS (50:50)
POPS
[A289C]
[R199Q]
[H230Q]
[K233Q]
[R291Q]

**Supplementary Fig. 11. Pharmacological activity and thermal melting behavior of $A_{2A}$AR variants in lipid nanodiscs.** Homologous competition binding experiments with the antagonist ZM241385 are shown in the left panels, and competition binding experiments with the agonist NECA are shown in the right panels for **a** $A_{2A}$AR[R199Q,A289C], **b** $A_{2A}$AR[H230Q,A289C], **c** $A_{2A}$AR[K233Q,A289C] and **d** $A_{2A}$AR[R291Q,A289C] in lipid nanodiscs containing POPC and POPS at a molar ratio of 70:30. The measured $K_D$ or $K_I$ values are shown in each panel. Error bars indicate the s.e.m for 3 independent trials. **c** Melting temperature (Tm) values calculated from fluorescence thermal shift assays for $A_{2A}$AR[A289C] and the variants $A_{2A}$AR[R291Q, A289C], $A_{2A}$AR[K233Q, A289C], $A_{2A}$AR[H230Q, A289C] and $A_{2A}$AR[R291Q, A289C] in nanodiscs containing 100% POPC (black bars), POPC and POPS at a 50:50 molar ratio (grey bars) and 100% POPS (white bars). Error bars for each sample were calculates as the standard error of the mean (s.e.m.) for n>3 independent experiments.

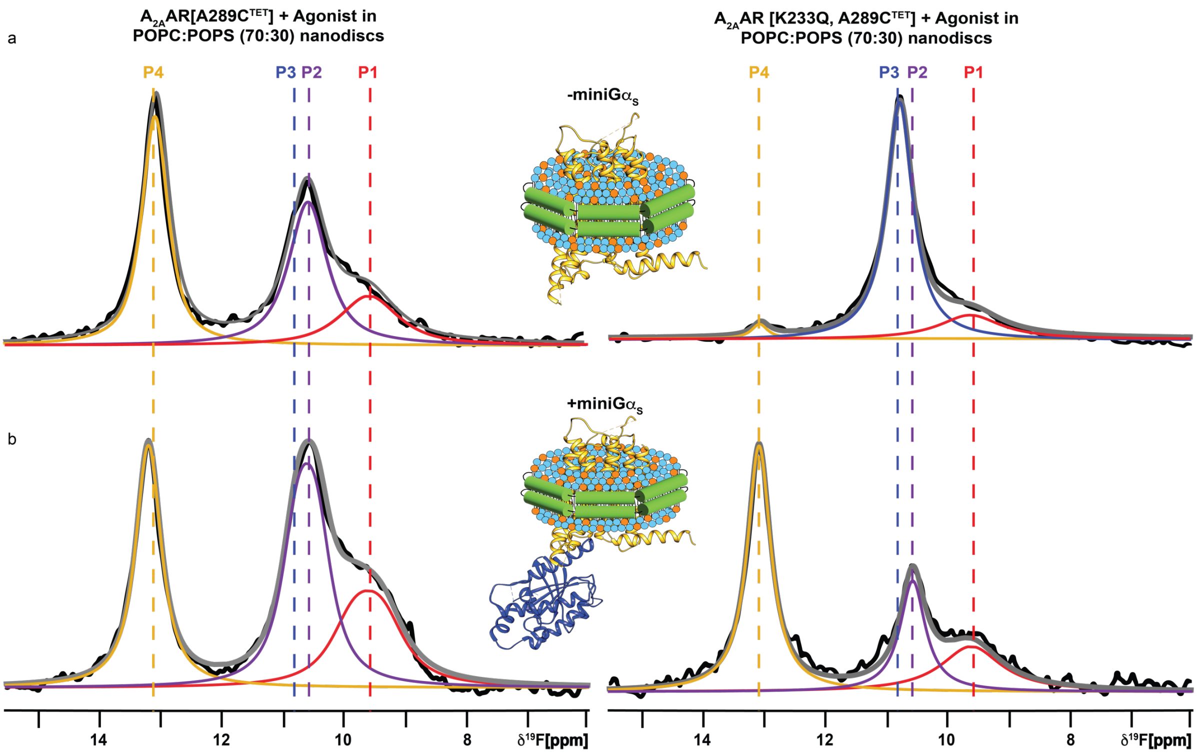


**Supplementary Fig. 12. $^{19}$F-NMR spectra of $A_{2A}$AR[K233Q,A289C$^{TET}$] complexes with the agonist NECA and NECA and mini-G$\alpha_S$ in nanodiscs containing mixtures of POPC and POPS.** $^{19}$F-NMR spectra of the $A_{2A}$AR[K233Q, A289C$^{TET}$] complex with **a** the agonist NECA and **b** NECA and mini-G$\alpha_S$ in nanodiscs containing POPC and POPS (70:30 ratio). Same presentation details as in Figure 1.

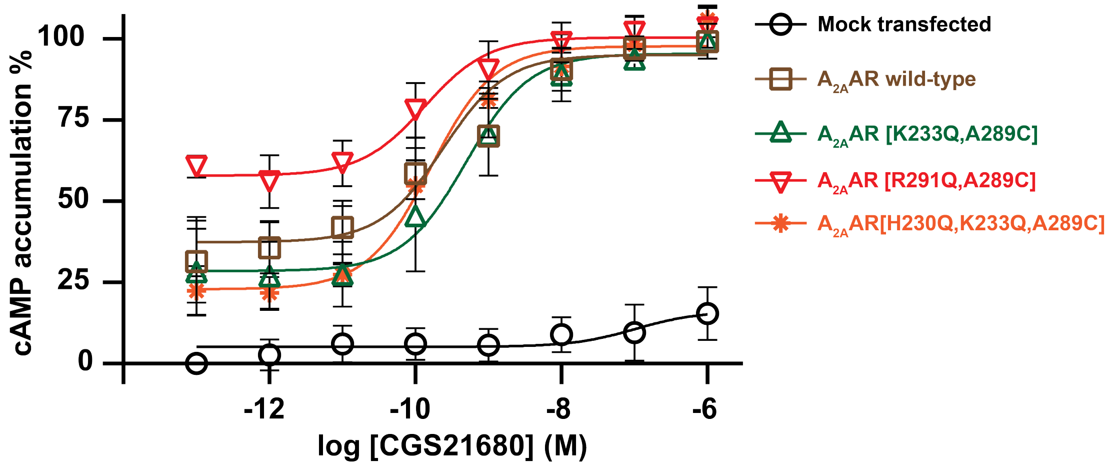


**Supplementary Fig. 13. Cell signalling assays with $A_{2A}$AR and $A_{2A}$AR variants.** cAMP accumulation experiments upon stimulation with the agonist CGS21680 for $A_{2A}$AR and several $A_{2A}$AR variant proteins. Data are shown as the means ± standard deviation. Experiments were conducted three times, each in triplicate.

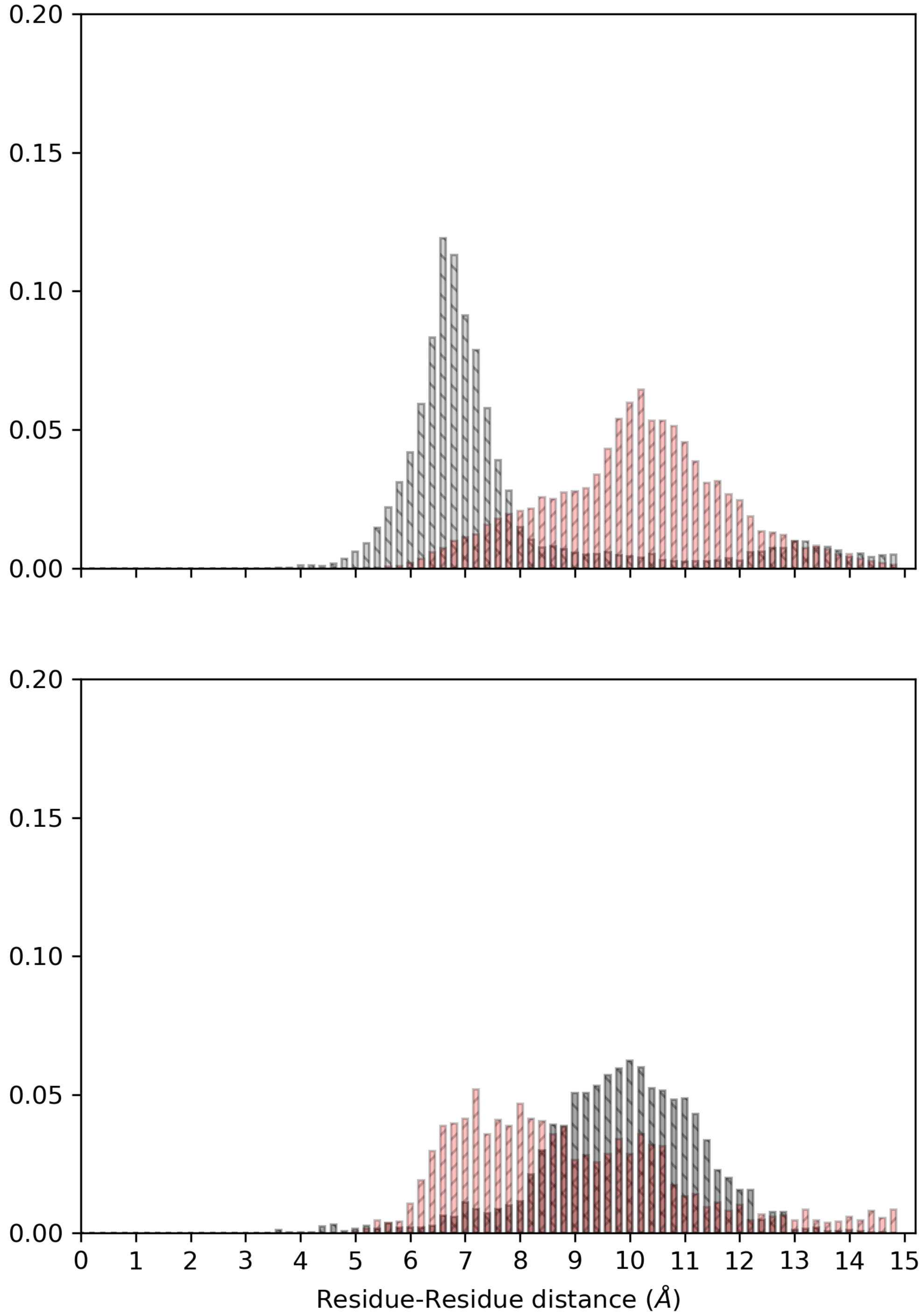
0.20
0.15
0.10
0.05
0.00
0 1 2 3 4 5 6 7 8 9 10 11 12 13 14 15
Residue-Residue distance (Å)

**Supplementary Fig. 14. Distance distribution between K233$^{6.35}$ and R291$^{7.56}$ in $A_{2A}$AR inactive and active conformers.** Top panel: Distance distribution between K233$^{6.35}$ and R291$^{7.56}$ in simulations of the antagonist-bound inactive state (black) and the fully active, G protein-coupled state (red). Bottom panel: distance distribution between K233$^{6.35}$ and R291$^{7.56}$ in simulations of the active state but in the absence of both POPS lipids and G protein (black), and the same receptor state in the presence of membranes with POPS (red).

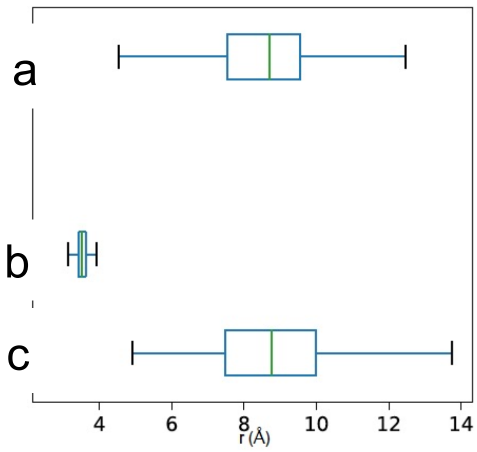


**Supplementary Fig. 15. Distance distribution between H230$^{6.32}$ in $A_{2A}$AR and E392 in the mini-G$\alpha_S$ protein.** The distance distribution between H230$^{6.32}$ and E392 on the mini-G$\alpha_S$ protein (a) is mimicked by the carboxylic acid headgroup of a PS if the receptor is simulated in the fully active state but without G-protein (c). However, if the receptor is instead simulated in the inactive state in the presence of PS (b), the H230$^{6.32}$ to PS distance is very narrow and short-ranged, very unlike the H230$^{6.32}$ and E392 distance. Vertical lines are the medians, boxes are the upper and lower quartile, and the horizontal lines are the total range.

**Supplementary Table 1.** Relative populations of $A_{2A}AR$ conformational states observed in $^{19}F$-NMR spectra

| Sample | Relative Populations[1] | | | |
|---|---|---|---|---|
| | P4 | P3 | P2 | P1 |
| **Samples prepared in DDM/CHS micelles** | | | | |
| $A_{2A}$AR[A289C]-NECA | 0.13 | 0.00 | 0.31 | 0.56 |
| $A_{2A}$AR[A289C]-ZM241385 | 0.00 | 0.54 | 0.00 | 0.46 |
| **Samples prepared in lipid nanodiscs** | | | | |
| $A_{2A}$AR[A289C]-NECA with POPC | 0.00 | 0.70 | 0.13 | 0.17 |
| $A_{2A}$AR[A289C]-NECA + miniG$\alpha_S$ with POPC | 0.18 | 0.00 | 0.50 | 0.32 |
| $A_{2A}$AR[A289C]-NECA with POPC:POPS (70:30) | 0.43 | 0.00 | 0.39 | 0.17 |
| $A_{2A}$AR[A289C]-ZM241385 with POPC:POPS (70:30) | 0.00 | 0.71 | 0.00 | 0.29 |
| $A_{2A}$AR[A289C]-NECA with POPC:POPA (70:30) | 0.48 | 0.00 | 0.29 | 0.23 |
| $A_{2A}$AR[A289C]-NECA with POPC:POPG (70:30) | 0.41 | 0.00 | 0.37 | 0.22 |
| $A_{2A}$AR[A289C]-NECA with POPC:PI(4,5)P2 (95:5) | 0.58 | 0.00 | 0.29 | 0.12 |
| $A_{2A}$AR[K233Q,A289C]-NECA with POPC | 0.22 | 0.00 | 0.41 | 0.37 |
| $A_{2A}$AR[H230Q,A289C]-NECA with POPC | 0.22 | 0.00 | 0.28 | 0.50 |
| $A_{2A}$AR[R291Q,A289C]-NECA with POPC | 0.02 | 0.73 | 0.00 | 0.25 |
| $A_{2A}$AR[K233Q,A289C]-NECA with POPC:POPS (70:30) | 0.03 | 0.79 | 0.00 | 0.18 |
| $A_{2A}$AR[H230Q,A289C]-NECA with POPC:POPS (70:30) | 0.00 | 0.66 | 0.00 | 0.34 |
| $A_{2A}$AR[R291Q,A289C]-NECA with POPC:POPS (70:30) | 0.58 | 0.00 | 0.11 | 0.32 |
| $A_{2A}$AR[K233Q,A289C]-NECA +miniG$\alpha_S$ with POPC:POPS (70:30) | 0.51 | 0.00 | 0.25 | 0.24 |

[1] Values are reported as a fraction of the total integral of all signals from 7.5 ppm to 14.5 ppm.

**Supplementary Table 2.** Sequences of oligonucleotides used to generate $A_{2A}$AR variants.

| Mutation | Directionality | Primer |
|---|---|---|
| A289C | Forward | TCGGTTGTGAATCCCTTCATTTACTGCTACCGTATCCGCGAGTTCCGCCAG |
| | Reverse | CTGGCGGAACTCGCGGATACGGTAGCAGTAAATGAAGGGATTCACAACCGA |
| R199Q | Forward | CTCATGCTGGGTGTCTATTTGCAAATCTTCCTGGCGGCGCGACGACAGCT |
| | Reverse | AGCTGTCGTCGCGCCGCCAGGAAGATTTGCAAATAGACACCCAGCATGAG |
| H230Q | Forward | TGCAGAAGGAGGTCCAAGCTGCCAAGTCACTGGCCATCATTGTGGGGCTC |
| | Reverse | GAGCCCCACAATGATGGCCAGTGACTTGGCAGCTTGGACCTCCTTCTGCA |
| K233Q | Forward | TGCAGAAGGAGGTCCATGCTGCCCAATCACTGGCCATCATTGTGGGGCTC |
| | Reverse | GAGCCCCACAATGATGGCCAGTGATTGGGCAGCATGGACCTCCTTCTGCA |
| R291Q | Forward | GTTGTGAATCCCTTCATTTACTGTTACCAAATCCGCGAGTTCCGCCAGAC |
| | Reverse | GTCTGGCGGAACTCGCGGATTTGGTAACAGTAAATGAAGGGATTCACAAC |

**Supplementary Table 3.** Details of systems simulated.

| Receptor State | Lipid Composition (PC/PS) | Replicas x Duration | Ionic Lock Distance [Å][1] |
|---|---|---|---|
| Inactive[2] | 85/15 | 5x500 nsec | 4.1 (0.5) |
| Inactive | 15/85 | 5x500 nsec | 4.2 (0.1) |
| Active w/ mini-G$\alpha$s[3] | 85/15 | 5x500 nsec | 19.7 (0.3) |
| Active w/ mini-G$\alpha$s | 15/85 | 5x500 nsec | 19.9 (0.5) |
| Restrained Active w/o mini-G$\alpha$s[4] | 100/0 | 5x500 nsec | 21.0 (0.7) |
| Restrained Active w/o mini-G$\alpha$s | 15/85 | 5x500 nsec | 21.0 (1.0) |
| Unrestrained active w/o mini-G$\alpha$s[5] | 15/85 | 5x400 nsec | 20.6 (0.9) |

[1] Distance between the sidechain nitrogens of Arg102$^{3.50}$ and oxygens of Glu228$^{6.30}$, reporting on the conformation of the ionic lock (D/ERY motif)

[2] Simulations initiated from the $A_{2A}$AR antagonist-bound structure PDB 3EML

[3] Simulations initiated from the $A_{2A}$AR ternary complex structure with agonist and mini-G$\alpha$s PDB 5G53

[4] Simulations initiated from the $A_{2A}$AR ternary complex structure with agonist and mini-G$\alpha$s PDB 5G53 where the mini-G$\alpha$s protein was deleted and the receptor backbone restrained

[5] Simulations initiated from the $A_{2A}$AR ternary complex structure with agonist and mini-G$\alpha$s PDB 5G53 where the mini-G$\alpha$s protein was deleted and simulations were allowed to continue for several hundred nanoseconds without backbone restraints
